# Unsupervised domain adaptation for the automated segmentation of neuroanatomy in MRI: a deep learning approach

**DOI:** 10.1101/845537

**Authors:** Philip Novosad, Vladimir Fonov, D. Louis Collins

**Author notes:** We would like to acknowledge funding from the Famille Louise and André Charron. This work was also supported in part by a doctoral fellowship from the Fonds de recherche du Québec – Santé (FRQS) and a grant from the Natural Sciences and Engineering Research Council of Canada (NSERC) CREATE (4140438 - 2012). P. Novosad is with the Neuroimaging and Surgery Technologies (NIST) group at McGill University, Montreal, Quebec, Canada V. Fonov is with the Neuroimaging and Surgery Technologies (NIST) group at McGill University, Montreal, Quebec, Canada D. L. Collins is with the Neuroimaging and Surgery Technologies (NIST) group at McGill University, Montreal, Quebec, Canada.

## Abstract

Neuroanatomical segmentation in T1-weighted magnetic resonance imaging of the brain is a prerequisite for quantitative morphological measurements, as well as an essential element in general pre-processing pipelines. While recent fully automated segmentation methods based on convolutional neural networks have shown great potential, these methods nonetheless suffer from severe performance degradation when there are mismatches between training (source) and testing (target) domains (e.g. due to different scanner acquisition protocols or due to anatomical differences in the respective populations under study). This work introduces a new method for unsupervised domain adaptation which improves performance in challenging cross-domain applications without requiring any additional annotations on the target domain. Using a previously validated state-of-the-art segmentation method based on a context-augmented convolutional neural network, we first demonstrate that networks with better domain generalizability can be trained using extensive data augmentation with label-preserving transformations which mimic differences between domains. Second, we incorporate unlabelled target domain samples into training using a self-ensembling approach, demonstrating further performance gains, and further diminishing the performance gap in comparison to fully-supervised training on the target domain.

## 1 Introduction

Structural segmentation in T1-weighted (T1w) magnetic resonance imaging (MRI) is a prerequisite for volume, shape, and thickness measurements, as well as an essential element in general pre-processing pipelines. While manual or semi-automatic labellings produced by trained human experts are widely considered the ‘gold standard’ approach for segmentation, such labellings are highly time-consuming and subject to both inter- and intra-rater variability. Consequently, much research has been devoted to developing fully automated methods for accurate and robust neuroanatomical segmentation. Recently, applications of convolutional neural networks (CNNs) [1] to the task of neuroanatomical segmentation have produced new state-of-the-art results [2, 3, 4, 5]. Despite these recent successes, such tools are commonly developed and validated under an overly restrictive assumption that both the training and testing data are sampled from the same underlying distribution (or ‘domain’). In T1w MRI, distributional shifts across domains are commonly observed due to variations in pulse sequences and scanner hardware, in addition to anatomical differences dependent on the imaged population. In practice, acquiring representative and high-quality labelled training data for each ‘target’ domain of interest is often infeasible, and pre-labelled training data from another ‘source’ domain are used for training instead. This domain mismatch can cause severe performance degradation, reducing the accuracy of subsequent analyses.

While some studies have advocated for semi-supervised approaches, whereby a network pre-trained on one or several source domains is fine-tuned on limited quantities of labelled target domain data (so-called ‘transfer learning’) [6], fully unsupervised approaches which do not require any manual labellings on the target domain are more desirable. Example CNN-based unsupervised domain adaptation approaches in medical imaging segmentation include that of Kamnitsas et al. [7], which adopted a domain-adversarial method for brain tumour segmentation in MRI, and that of Perone et al. [8], which adopted a self-ensembling method for spinal cord segmentation in MRI. To the best of our knowledge, no work has specifically addressed the domain adaptation problem for general neuroanatomical segmentation, particularly in highly challenging scenarios where source and target domains differ not only with respect to scanner acquisition protocol (resulting in differences with respect to overall image brightness, contrast, noise and resolution), but also with respect to anatomy (e.g. due to differences in age and/or health) of the brains of the scanned individuals.

In this work, we propose an extension to our previously developed CNN-based method for neuroantomical segmentation [2] which is specifically designed for fully unsupervised domain adaptation in challenging T1-weighted (T1w) neuroanatomical segmentation applications. First, we demonstrate that networks with greater domain generaliz-ability can be trained using an appropriate data augmentation scheme with random transformations designed to mimic inter-domain differences in T1w MRI. Second, we incorporate unlabelled target domain samples into training using a self-ensembling [9, 10, 11] approach. Using three different manually annotated datasets, we extensively validate our method and compare it with a domain-adversarial [12, 7] approach for unsupervised domain adaptation, as well as a classic patch-based [13] segmentation approach, in each case demonstrating improved cross-domain performance.

## 2 Methods and materials

### 2.1 Unsupervised domain adaptation

During training, we assume that we have access to training samples ***x***^*S*^ and ***x***^*T*^ from the source and target domains respectively. However, only for samples from the source domain ***x***^*S*^ are the corresponding reference labels ***y***^*S*^ known. If the source domain and target domain are sufficiently similar, then a labeller network trained on labelled source samples can be simply applied to samples from the target domain. Unfortunately, the transferability of features learned by deep neural networks is limited due to fragile co-adaptation and representation specificity [14], leading to suboptimal performance on target domain samples in many cases. The task of unsupervised domain adaptation is to remedy this problem, i.e. to learn a labeller network *L*(***x***, ***θ***): *X → Y* which accurately predicts labels ŷ^*T*^ for inputs *x*^*T*^ from the target domain, i.e. which is adapted to the target domain.

A popular class of methods for domain adaptation applied to convolutional neural networks addresses this problem by seeking a labeller network for which the classification accuracy is high on labelled source samples, while simultaneously generating similar feature distributions across domains [15, 16, 12]. Methods belonging to this class differ primarily with respect to the specific choice of representation space in which to measure the disparity between domains (e.g. which network layer(s) to examine for inter-domain differences), and the choice of how to measure and minimize the distance. For example, the work of Ganin et al. [12] (extended to tumour-based segmentation in Kamnitsas et al. [7]) uses a domain-adversarial approach in which a classifier network is trained to simultaneously minimize the classification loss on labelled source samples while countering a domain-discriminator network in order to generate domain-invariant deep features. This approach can be too restrictive in cases where there is reason to expect that the distributions of feature maps should not be particularly similar across domains (e.g. if the two domains differ with respect to overall anatomy). Less restrictive approaches, which aim to match only lower-order statistics of deep feature distributions between domains have also been proposed [17, 18]. Nonetheless, even if distributions of the source and target deep features can be well aligned, there is no guarantee that the aligned target samples will fall on the correct sides of the learned decision boundary.

### 2.2 Self-ensembling for domain adaptation

A related approach for semi-supervised learning, called ‘self-ensembling’ [10, 11], incorporates unlabelled samples into training using an auxiliary *consistency loss*. The consistency loss penalizes differences between outputs of the network evaluated on the same input but under different *label-preserving* data augmentation transformations. Minimizing the consistency loss therefore helps to construct a regularized model which produces smoothly varying outputs with respect to it’s input, i.e. which is smooth around the (labelled and unlabelled) training data. This approach can also be interpreted as extrapolating the labels for the unlabelled samples [19], akin to so-called ‘label-propagation’ methods [20].

Self-ensembling has been recently extended to domain adaptation by French et al. [9], demonstrating state-of-the-art results for digit classification tasks, and applied to the task of domain adaptation for spinal cord grey matter segmentation in MRI by Perone et al [8]. As argued by French et al., since self-ensembling works by label propagation, it is crucial that the source and target domains at least partially overlap in input space. To encourage sufficient overlap between domains, the same authors propose an extensive set of label-preserving data augmentation transformations tailored to their particular task of digit recognition. In our work, we propose a set of label-preserving data augmentation transformations better suited for domain adaptation in T1-weighted MRI, which we describe in section II-D. Also as suggested by French et al., we maintain an exponential moving average (EMA) of the network parameters ***θ*** during training:

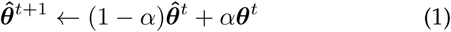

where *t* is the training batch, ***θ***^*t*^ are the network parameters at training batch *t*, *α* controls the ‘memory’ of the EMA (e.g. smaller values of *α* discount older observations faster) and 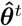 is the EMA of the network parameters at training batch *t*. Rather than comparing the output of the same network for two randomly transformed versions of the same input, we compare the output of the ‘student’ model (the network with parameters ***θ***) with that of the ‘teacher’ model (the same network but with the EMA parameters 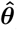). This approach has the benefit of encouraging the student model to more closely mimic the teacher model (which will tend to be a more accurate model [21]), in turn producing a beneficial feedback loop between the student and the teacher models [10].

We now explicitly formulate our method and training strategy for domain adaptation based on self-ensembling (Fig. 1). We first assume that we have access to a set of *N* samples from each of the source and target domains. The source loss 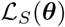 is computed over labelled source samples only as

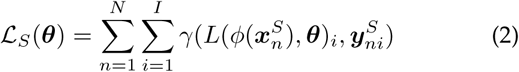

where *γ*(⋅) is the categorical cross-entropy function, *φ*(⋅) applies a random label-preserving data augmentation transformation to its input, 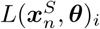 is the softmax output containing the predicted class probabilities at pixel *i*, and 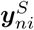 is the one-hot encoded reference label for input 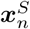. The target consistency loss 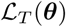 is computed over unlabelled target samples as

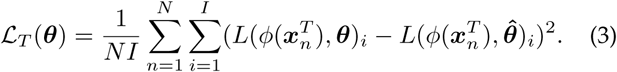

**Fig. 1.**
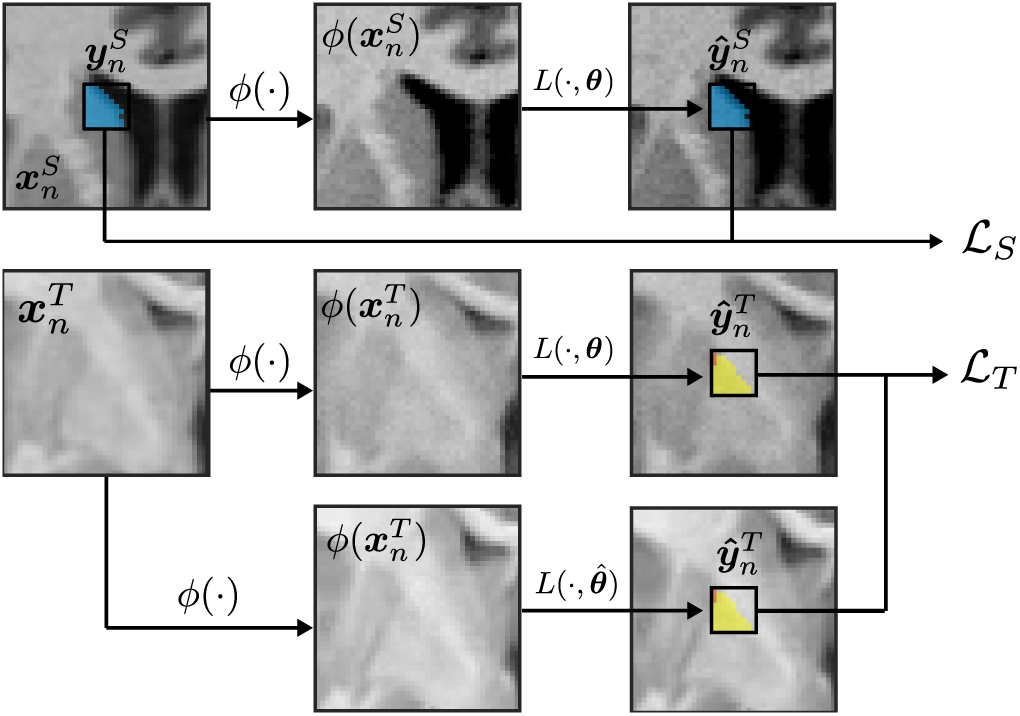
Self-ensembling for domain adaptation. Labelled source samples (top row) are used to maximize the labelling accuracy. In parallel, unlabelled target domain samples (bottom two rows) are used to minimize a consistency loss which penalizes differences between label predictions made on two randomly transformed versions of the same input.

The total loss function to minimize is given by

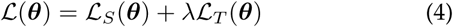

where *λ* is a hyperparameter which specifies the trade-off between accuracy on labelled source samples and target consistency.

### 2.3 Network architecture

We use the labeller network described in Novosad et al. [2], which combines a deep three-dimensional fully convolutional architecture with spatial priors. Spatial priors are incorporated by using a working volume to restrict the area in which samples are extracted (during both training and testing) and by explicitly augmenting the input with spatial coordinate patches. The network takes as input a large patch of size 41^3^×*N* (where *N* is the number of channels, including spatial coordinate patches) and first processes it using a series of sixteen 3 × 3 × 3 convolutional layers (applied without padding and with unary stride), reducing the size of the feature maps to 9^3^ (we note that each application of such a convolutional layer reduces the size of the feature maps by 1 voxel in each dimension). The output of each preceding convolutional layer is cropped and concatenated to produce a multi-scale representation of the input, which is further processed by a series of three 1 × 1 × 1 convolutional layers, producing a probabilistic local label estimate for the central 9^3^ voxels of the input for each of the *C* structures under consideration.

### 2.4 Increased domain generalizability using data augmentation

As shown in Fig. 2 and Fig. 3, T1w images from different datasets broadly differ with respect to both low-level (e.g. image brightness, contrast, resolution and noise) and high-level (anatomical) properties. Diversifying the appearance of training samples with respect to these properties can help train models which are more robust to differences among them. To this end, we use extensive data augmentation scheme consisting of random label-preserving (as required for compatibility with self-ensembling) transformations. Specifically, we explore five task-specific data augmentation techniques intended to increase inter-domain generalizability in T1w MRI, which we now describe. We additionally note that prior to applying the data augmentation transformations, images are pre-processed to zero mean and unit standard deviation as described in Section II-F.

**Fig. 2.**
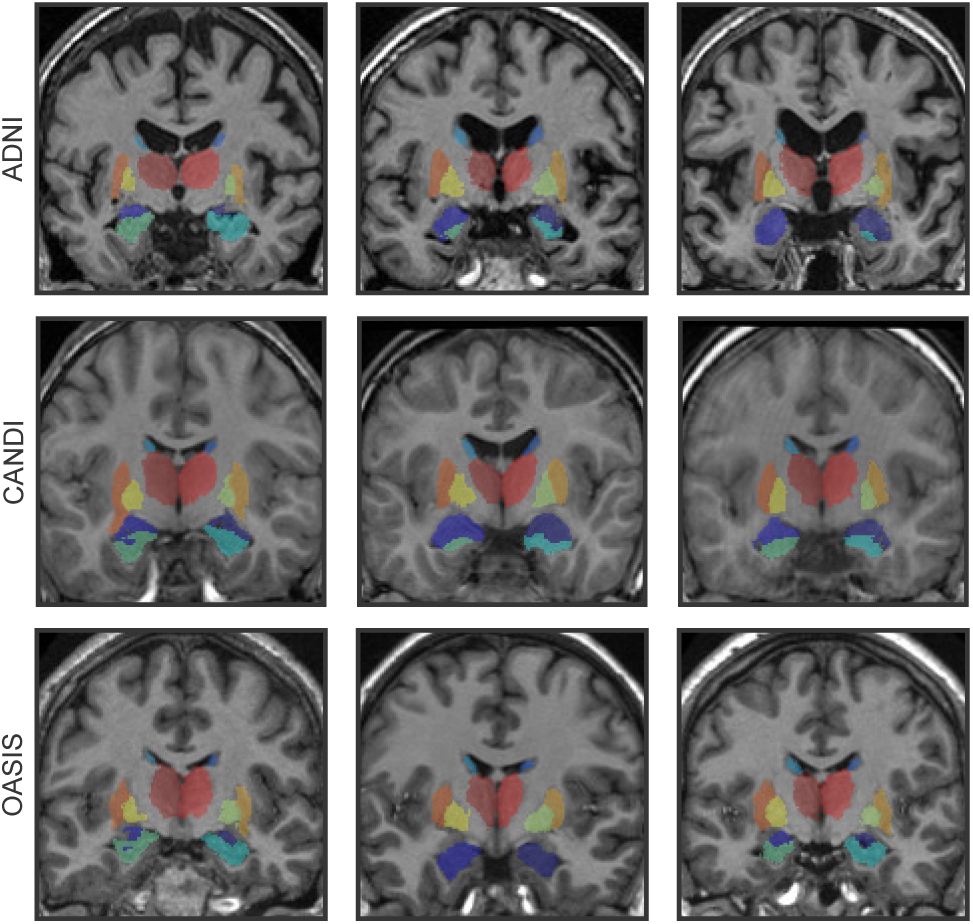
Three different pre-processed T1w datasets or ‘domains’ used in this work. Images from the different domains differ with respect to both low-level properties (e.g. image brightness, contrast, resolution and noise) and high-level anatomical properties.

**Fig. 3.**
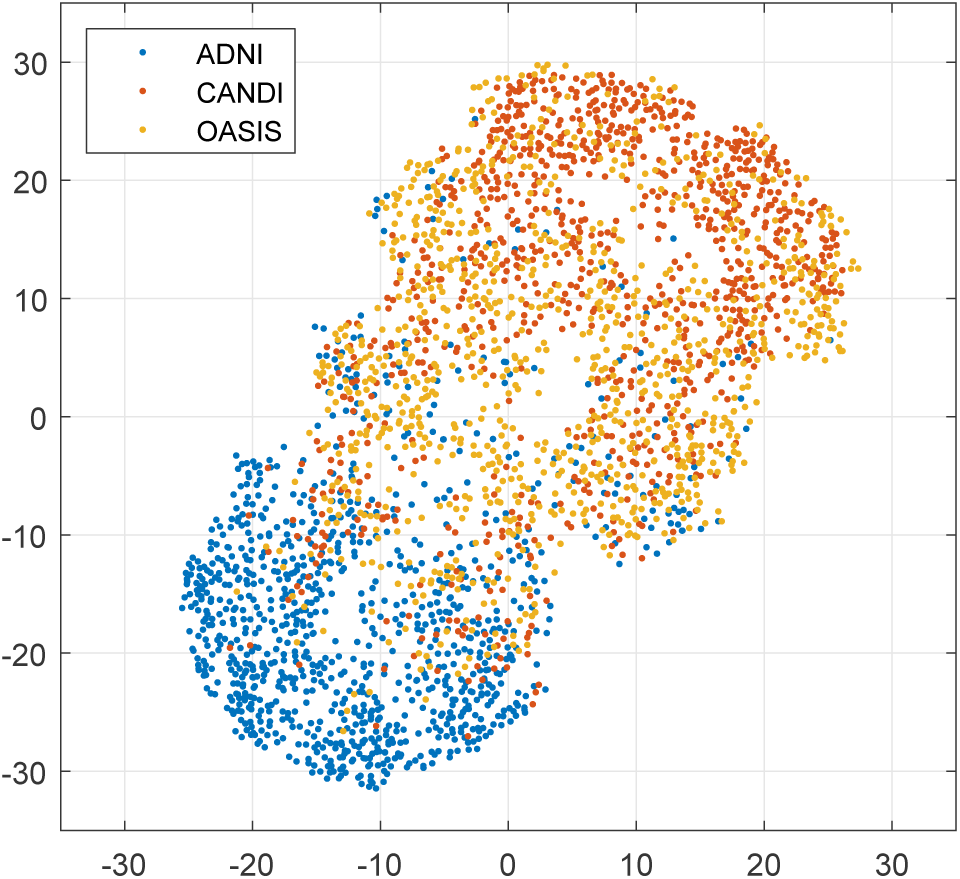
A two-dimensional non-linear embedding using t-SNE shows that samples from the different domains shown in Fig. 2 (here, input samples to the CNN, of size 41^3^) occupy different but overlapping regions of the input space.

1. *Brightness*: a random uniform offset is added to the sample:

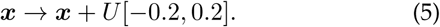
2. *Contrast*: the mean separation between low- and high-intensity voxels in the sample is randomly altered:

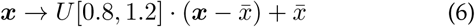

where 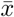 is the mean value of the sample ***x*** over all voxels.
3. *Sharpness*: high-frequency detail is randomly enhanced or suppressed:

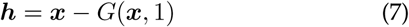

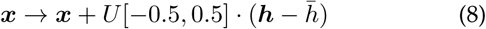

where ***h*** is a high-frequency image obtained by subtracting a Gaussian blurred (with standard deviation of 1) version of the sample from itself.
4. *Noise*: independent random Gaussian noise with zero mean and standard deviation 0.05 is added to each voxel of the sample:

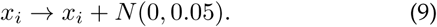
5. *Spatial deformations*: a random elastic deformation is applied to the sample. We use the approach described in [2] to generate the deformation fields with previously validated parameters *σ* = 4 mm and *α* = 2 mm. As required for self-ensembling, however, the random deformation must be label-preserving (i.e. the labels of the central 9^3^ voxels of each original sample should remain consistent with its transformed variant). To this end, we create a binary mask image with the same spatial dimensions as the training sample, and set the central (9 + 3*σ*)^3^ voxels to 0 and the remaining voxels to 1. We then blur the mask image with a Gaussian filter with standard deviation *σ*, such that the value of the blurred mask is approximately zero for the central 9^3^ voxels, and then smoothly increasing to one at the edges. In this way, the deformation randomly warps the background anatomy while preserving the labels of the original sample. We note that the parameters associated with each transformation were selected heuristically in order to produce random samples with realistic appearances. Example randomly transformed samples are displayed in Fig. 4.

**Fig. 4.**
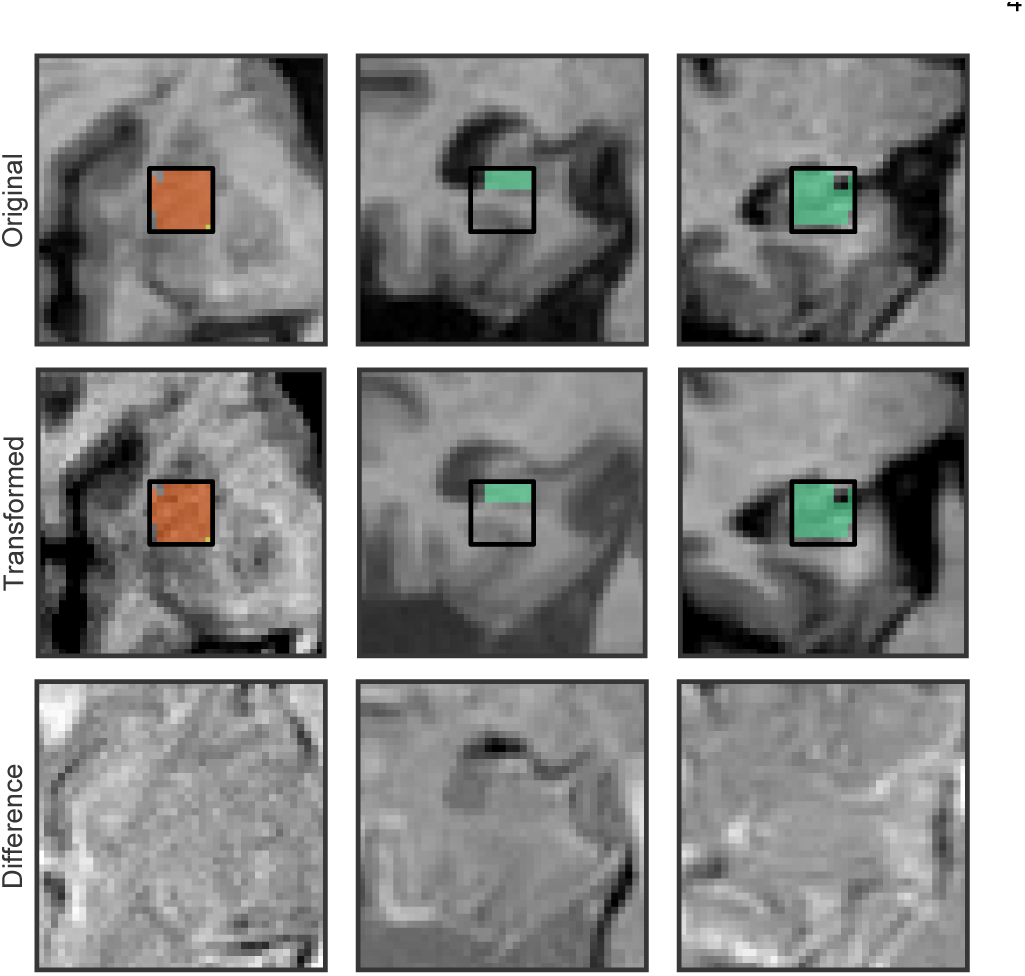
Example data augmentation transformations applied to samples from the ADNI dataset. The original samples are displayed in the first row, and the transformed versions in the second row. The respective intensity differences between the original and transformed samples are displayed in the third row. Here, each of the five transformations described in section II-D are applied to each sample in a random order. Note that the transformations do not alter the label of the central 9^3^ voxels (small box), which is essential for the target consistency loss (Equation (3)).

### 2.5 Training and testing

Training and testing for the non-adapted networks is done as described in [2]. For the proposed self-ensembling approach, a number of modifications were required for training, which are now discussed in turn.

#### 2.5.1 Pseudo-labelling for approximate class balancing

Using approximately class-balanced training samples is required to ensure that the learned networks are not biased against smaller structures. In [2], training samples are drawn such that the central voxel is equally likely to belong to any of the structures under consideration. Since no reference labels are available on the target domain, we instead use a model pre-trained on the source domain to generate pseudo-labels, which are in turn used during training to extract (approximately) class balanced samples from the target domain.

#### 2.5.2 Fine-tuning

Rather than training the domain-adapted network from scratch, we opt to fine-tune the network pre-trained on the source domain only. In our preliminary studies, we found that this approach resulted in faster and more stable training, allowing us to drop the ‘ramp-up’ term (used in the works of French et al. [9] and Perone et al. [8]) required to slowly increase the consistency loss in order to stabilize training.

#### 2.5.3 Batch normalization statistics

As done in the work of French et al. [9], we compute the loss in equation (4) at each iteration by passing through two separate batches: one batch of labelled source-domain samples (computing the supervised classification loss) and one batch of unlabelled target-domain samples (computing the unsupervised consistency loss), and then form a weighted sum before backpropagating the loss to update the network parameters. In the work of [9], inspired by [17], the authors opt for an approach whereby the source and target samples are batch-normalized independently during training. While this approach ensures that the network produces feature maps with similar mean and variance regardless of the input domain, it does not ensure that the same classes across domains are mapped to similar features. Indeed, in the presence of strong anatomical differences between domains, this approach can directly cause such a discrepancy. In our implementation of self-ensembling we instead use consistent batch normalization statistics for both domains: when fine-tuning the pre-trained source network using self-ensembling, we freeze the batch normalization layers and instead use the pre-computed running average batch statistics from the source domain during both training and testing.

#### 2.5.4 Training specifications

Network parameters are optimized iteratively using RM-SProp [22], an adaptive stochastic gradient descent algorithm with Nesterov momentum [23] (momentum = 0.9) for acceleration. At each epoch, we sample approximately 1500 voxels from the images in the source and target domains, with an equal number of voxels sampled from each training subject in each domain, and such that an equal number of voxels are extracted from each structure (background included). Training samples (i.e. whole patches with spatial coordinates [2]) are then extracted around each selected voxel. The samples are then processed iteratively in mini-batches of size 16. We maintain the exponential moving average network for the self-ensembling method using *α* = 0.99 following recommendations by Tarvainen et al. [10]. We additionally regularized the network using the *L*_2_ norm on the weights with regularization weight set to 10^*−*4^.

Network weights are randomly initialized with the Glorot method [24] and all biases are initialized to zero. A static learning rate of 1×10^*−*4^ was used. Because no validation set is available on the target domain to drive early-stopping, the networks were trained for a fixed number of 50 epochs. We note that preliminary experiments showed little improvement in the unsupervised loss after this point.

Training was performed on a single NVIDIA TITAN X with 12GB GPU memory. Software was coded in Python, and used Lasagne (https://lasagne.readthedocs.io/en/latest/index.html), a lightweight library to build and train the neural networks in Theano [25].

### 2.6 Preprocessing and Datasets

For validation, we use three different T1w datasets with labels provided by Neuromorphometrics (http://www.neuromorphometrics.com). Image preprocessing consisted of non-uniformity correction with the N3 algorithm [26], 12-parameter affine registration to the MNI-ICBM152 template using an in-house MINC (https://bic-mni.github.io/) registration tool based on normalized mutual information [27], and intensity normalization to zero mean and unit standard deviation. In our studies, we focus on segmentation of the hippocampus as well as the following subcortical structures and the left and right thalamus, caudate, putamen, pallidum, hippocampus and amygdala for a total of 13 classes (one class being background). Representative images from each dataset, after preprocessing, are displayed in Fig. 2 with labels overlaid. Dataset details are provided below.

#### 2.6.1 ADNI dataset

The Alzheimer’s Disease Neuroimaging Initiative (ADNI) dataset [28, 29] used in this work contains images of 30 subjects (minimum/mean/maximum age = 62.4/75.7/87.9 years), with 15 images from subjects with Alzheimer’s disease, and 15 images from healthy elderly subjects. These images were acquired on 1.5 T General Electric (GE), Philips, and Siemens scanners using a magnetization-prepared rapid acquisition gradient-echo (MP-RAGE) sequence.

#### 2.6.2 CANDI dataset

The Child and Adolescent NeuroDevelopment Initiative (CANDI) dataset [30] used in this work contains images of 13 young subjects, some of which have been diagnosed with psychiatric disorders (minimum/mean/maximum age = 5/9.5/15 years), acquired on a 1.5 T GE Signa scanner using an inversion recovery-prepared spoiled gradient recalled echo sequence.

#### 2.6.3 OASIS dataset

The Open Access Series of Imaging Studies (OASIS) dataset [31] used in this work contains images of 20 healthy young adults (minimum/mean/maximum age = 19/23.1/34 years) acquired on a Siemens 1.5 T Vision scanner using an MPRAGE sequence.

## 3 Experiments and Results

We assess segmentation accuracy using the Dice coefficient. The Dice coefficient measures the extent of spatial overlap between two binary images. The Dice coefficient is defined as 100%×2|*A ⋂ R| /*(|*A*| + |*R*|) where *A* is an automatically segmented label image, *R* is the reference label image, is the intersection, and|⋅| counts the number of non-zero elements. We here express the Dice coefficient as a percentage, with 100% indicating perfect overlap. We note that for multi-label images, we compute the Dice coefficient for each structure independently.

To reduce the variability in our performance estimate of the various CNN-based methods, we report mean performance estimates over multiple independent runs (10 runs for the results in section III-A (since individual runs were more variable) and 5 runs for the results in sections III-B and III-C), i.e. re-training and re-testing each network using different random seeds.

### 3.1 Effect of data augmentation

We assessed the effect of each data augmentation transformation described in section II-D, (brightness, contrast, sharpness, noise and spatial deformations) on domain generalizability by training networks on the OASIS subjects using each or all types of transformation (in the latter case, each transformation was applied to each sample in a random order), and then applying the networks to segment both ADNI and CANDI datasets. Mean Dice coefficients are reported in Table I.

**TABLE 1.** Impact of various data augmentation transformations on domain generalizability for the OASIS → ADNI and OASIS → ADNI experiments. Mean Dice coefficients (with standard deviation in parentheses) across 10 independent runs are reported.

| | OASIS → ADNI | OASIS → CANDI |
| --- | --- | --- |
| None | 71.1 (1.4) | 72.7 (1.0) |
| Brightness | 74.4 (1.3) | 75.6 (0.7) |
| Contrast | 71.8 (1.2) | 72.7 (1.3) |
| Noise | 71.2 (1.8) | 72.4 (1.8) |
| Sharpness | 73.1 (1.5) | 73.2 (0.9) |
| Deformations | 72.5 (1.3) | 72.8 (0.8) |
| All | 75.6 (1.1) | 76.6 (0.8) |

Brightness transformations were the most effective for both OASIS → ADNI and OASIS → CANDI (*p <* 6 × 10^*−*5^, paired t-test, compared to baseline without data augmentation), followed by sharpness transformations and random spatial deformations. Contrast transformations improved performance in the OASIS → ADNI adaptation, but the effect was not significant compared to the baseline (*p* = 0.25), and had no effect on the OASIS → CANDI adaptation. The effect of random noise addition did not significantly improve performance relative to the baseline in either case (*p >* 0.6). Finally, the combination of all five data augmentation transformations produced the best performance for both source → target tasks.

### 3.2 Effect of consistency loss

Next we assessed the impact of the parameter *λ* in equation (4), which controls the influence of the consistency loss on unlabelled target domain samples. Again we train networks using the OASIS dataset as the source domain, and consider both ADNI and CANDI as separate target domains. Mean Dice coefficients are plotted in Fig. 5. We note that here extensive data augmentation was included in all experiments using all five data augmentation transformations described in section II-D.

**Fig. 5.**
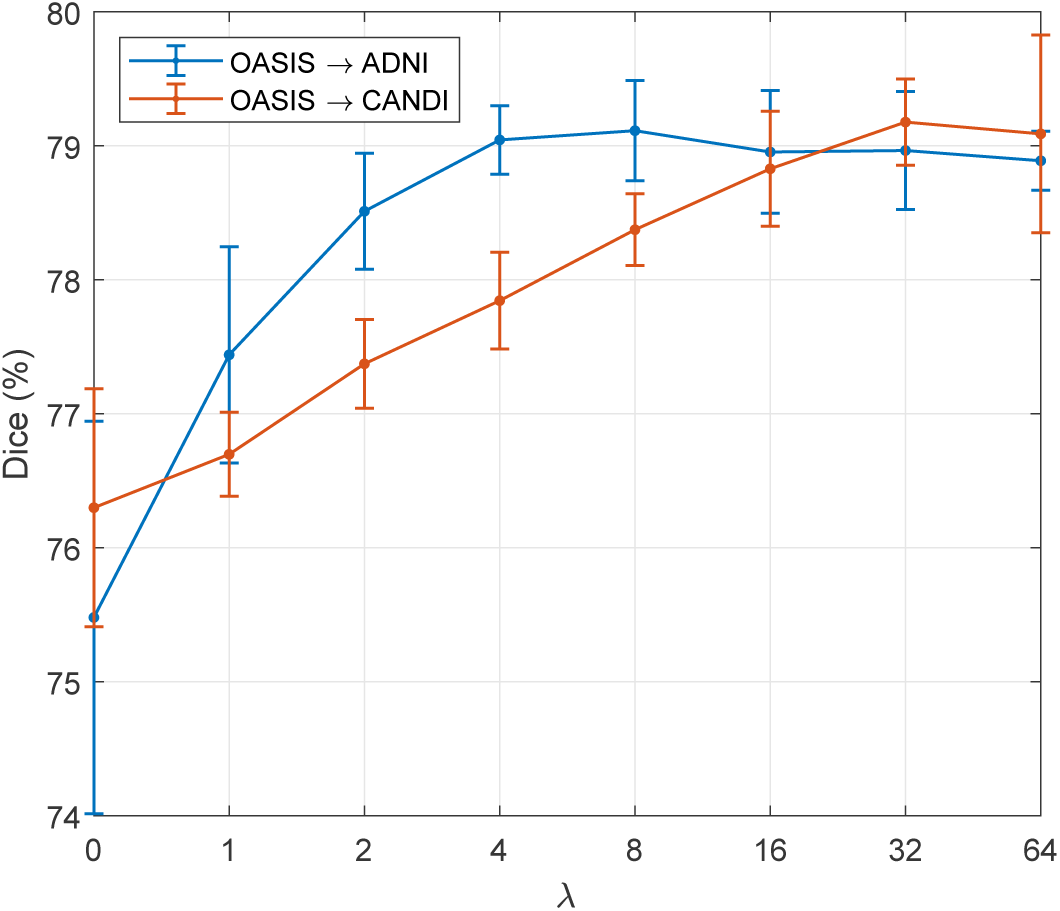
Impact of target consistency weight *λ* on self-ensembling performance in both OASIS → ADNI and OASIS → CANDI adaptations. Mean and standard deviation of Dice coefficients across all structures are shown over 5 independent runs. While performance is generally robust to the choice of *λ*, both adaptations achieved near optimal performance at *λ* = 32.

Performance in the OASIS → ADNI adaptation increased with increasing *λ* in the range [1, 8] and then reached a plateau, while performance in the OASIS → CANDI adaptation increased more slowly with increasing *λ* in the range [1, 32]. In general, the performance of self-ensembling was robust to the choice of *λ* for a wide range of values, and both adaptations achieved near optimal performance for *λ* = 32. We therefore use this value in the subsequent experiments, regardless of the specific source *→* target task

### 3.3 Comparison of methods

For comparison, we also consider our own implementation of the domain-adversarial (DA) method [12], previously applied to the task of domain adaptation in MR segmentation by Kamnitsas et al. [7]. This method minimizes the classification loss on labelled source samples while learning domain-invariant features by countering a domain-discriminator network which attempts to predict the domain of the input data by observing the generated features. In our implementation, we use the same labeller network as for self-ensembling (section II-C) and attach the domain-discriminator to the last layer (immediately prior to the final softmax activation function). The discriminator consisted of four 3 × 3 × 3 convolutional layers (applied without padding) each with 32 filters, and exponential linear units (ELUs) [32] were used as activation functions for all layers except the final one, which used a sigmoid function.

Table II reports mean Dice coefficients obtained by applying each method to each source → target adaptation. For comparison, we also provide results from fully supervised training (5-fold cross-validation) on the target domain, which can be interpreted as an upper-bound performance achievable by the domain adaptation methods. We first note that the addition of data augmentation improved performance of the baseline network in the inter-domain experiments as well as in intra-domain experiments. However, the improvement in the inter-domain experiments (increase in mean Dice, across all structures and all source → target adaptations, of 2.9%, from 73.2% to 76.1%) was considerably larger than in the latter experiments (increase in mean Dice of 1.2%, from 82.7% to 83.9%). This confirms that the data augmentation transformations used in this work are particularly effective at improving the domain generalizability of the trained networks. Indeed, the addition of data augmentation alone was more effective (*p <* × 10^*−*9^, paired t-test) than the domain-adversarial method, though less effective compared to the self-ensembling method using only minimal augmentation in the form of Gaussian noise. The combination of data augmentation and the domain-adversarial method produced a mean Dice coefficient of 77.3%, significantly better than either data augmentation or the domain-adversarial method alone (*p <* × 10^*−*9^), though comparably effective compared to self-ensembling with minimal augmentation (*p >* 0.05). Finally, the self-ensembling approach performed best of all unsupervised domain adaptation methods, producing a mean Dice coefficient of 78.4% (*p <* × 10^*−*9^ compared to the second best (domain-adversarial) method).

**TABLE 2.**
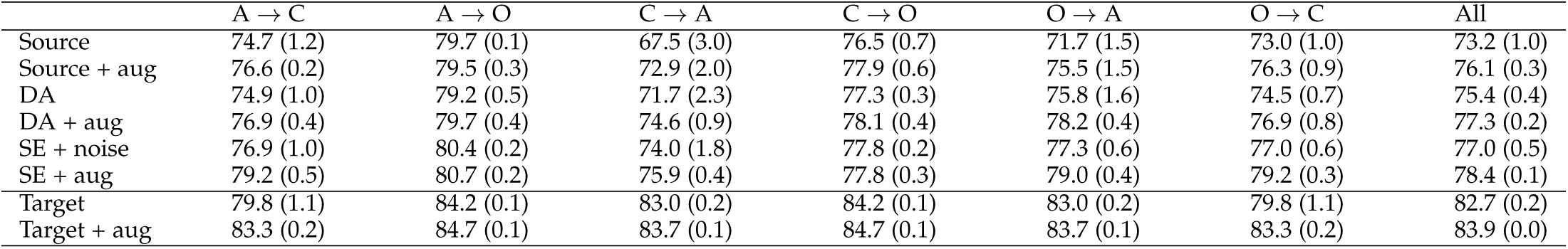
Comparison of segmentation methods (DA: domain-adversarial, SE: self-ensembling, aug: data augmentation) in all six source → target tasks. Mean Dice coefficients (with standard deviation in parentheses) across 5 independent runs are reported. The bottom two rows report performance across 5 independent runs of fully supervised training on the target domain, each using a 5-fold cross validation. A: ADNI, C: CANDI, O: OASIS.

| | A → C | A → O | C → A | C → O | O → A | O → C | All |
| --- | --- | --- | --- | --- | --- | --- | --- |
| Source | 74.7 (1.2) | 79.7 (0.1) | 67.5 (3.0) | 76.5 (0.7) | 71.7 (1.5) | 73.0 (1.0) | 73.2 (1.0) |
| Source + aug | 76.6 (0.2) | 79.5 (0.3) | 72.9 (2.0) | 77.9 (0.6) | 75.5 (1.5) | 76.3 (0.9) | 76.1 (0.3) |
| DA | 74.9 (1.0) | 79.2 (0.5) | 71.7 (2.3) | 77.3 (0.3) | 75.8 (1.6) | 74.5 (0.7) | 75.4 (0.4) |
| DA + aug | 76.9 (0.4) | 79.7 (0.4) | 74.6 (0.9) | 78.1 (0.4) | 78.2 (0.4) | 76.9 (0.8) | 77.3 (0.2) |
| SE + noise | 76.9 (1.0) | 80.4 (0.2) | 74.0 (1.8) | 77.8 (0.2) | 77.3 (0.6) | 77.0 (0.6) | 77.0 (0.5) |
| SE + aug | 79.2 (0.5) | 80.7 (0.2) | 75.9 (0.4) | 77.8 (0.3) | 79.0 (0.4) | 79.2 (0.3) | 78.4 (0.1) |
| Target | 79.8 (1.1) | 84.2 (0.1) | 83.0 (0.2) | 84.2 (0.1) | 83.0 (0.2) | 79.8 (1.1) | 82.7 (0.2) |
| Target + aug | 83.3 (0.2) | 84.7 (0.1) | 83.7 (0.1) | 84.7 (0.1) | 83.7 (0.1) | 83.3 (0.2) | 83.9 (0.0) |

We note that the performance of the baseline network (trained on the source domain only) was highly variable between independent runs (see standard deviations reported in Table II). This is expected, since the shape of the learned decision boundary is only constrained in the vicinity of the support of the source domain. The addition of data augmentation effectively increased the overlap of the transformed samples with target domain samples, reducing inter-run variability. Finally, the self-ensembling approach further reduced interrun variability to a level comparable to that of supervised training on the target domain.

Example segmentations are displayed in Fig. 6 and Fig. 7 for the OASIS *→* ADNI and OASIS *→* CANDI tasks, respectively. In the OASIS → ADNI application, erroneous segmentations produced by the baseline network (trained on the source domain only) were generally anatomically non-contiguous and characterized by large segments of missing labels (rows 1 and 2 of Fig. 6). In the OASIS CANDI application, erroneous segmentations produced by the same baseline network were generally characterized by anatomical irregularity (rows 1 and 2 of Fig. 7) and isolated clusters of spatially disconnected labels. In general, using the self-ensembling method with data augmentation minimized these errors, instead making errors concentrated along the more ambiguous structural boundaries.

**Fig. 6.**
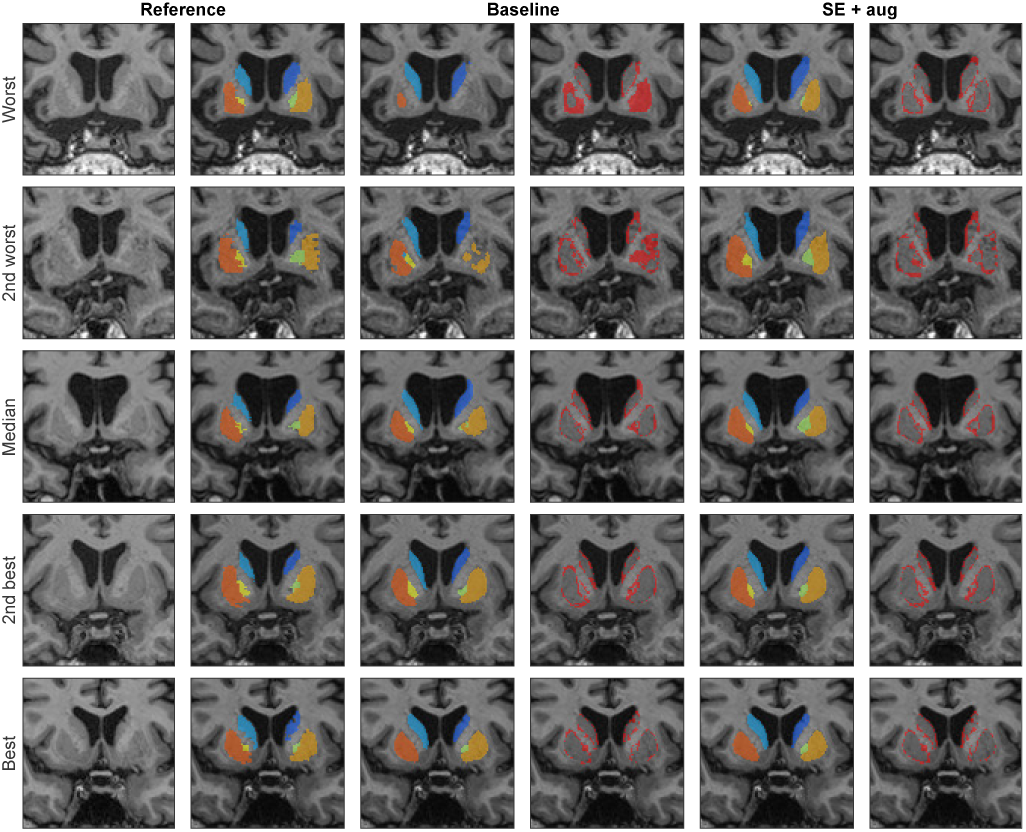
Example segmentations in the OASIS → ADNI task. The subjects with the worst, 2nd worst, median, 2nd best and best mean Dice coefficients after segmentation with the baseline network were chosen for comparison. Errors relative to the reference labels are shown in the fourth and sixth columns in red. Compared to the baseline network, the proposed segmentation method produced more anatomically contiguous segmentations consistent with the reference labels.

**Fig. 7.**
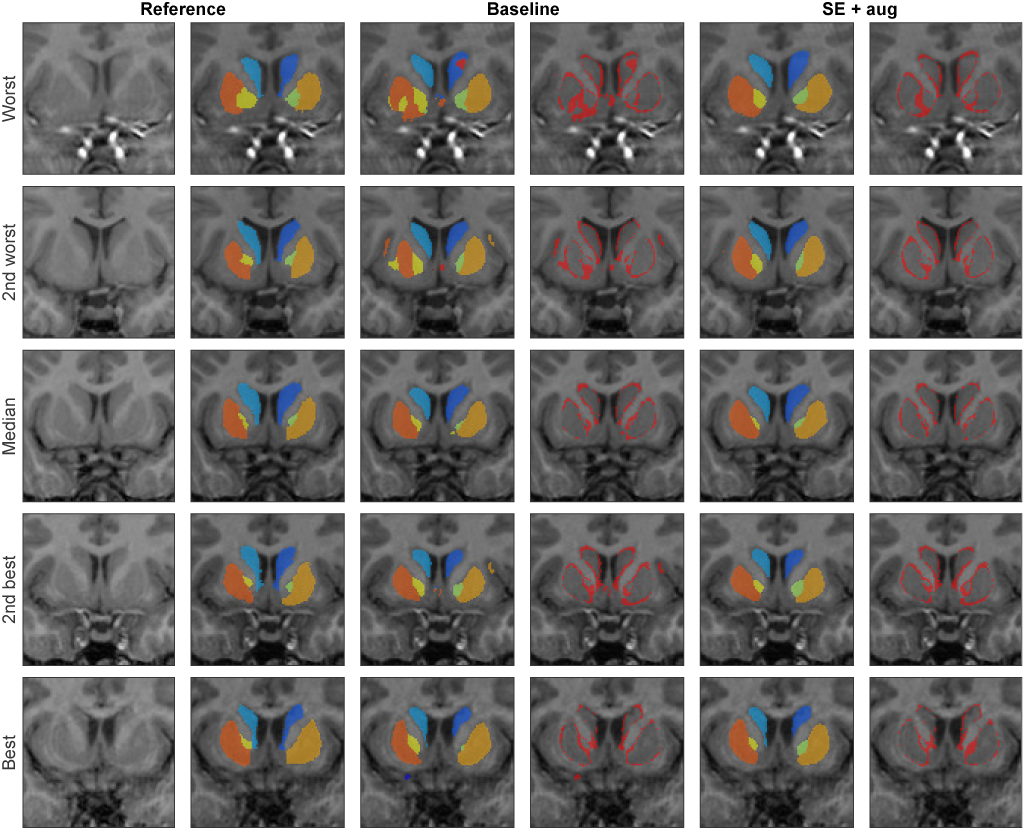
Similar to Fig. 6 but for the OASIS CANDI → task. While the baseline segmentation method produce anatomically irregular segmentations and isolated clusters of false positives (e.g. first two rows) these errors were avoided when using the proposed segmentation method.

For completeness, we also consider a classic patch-based segmentation (PBS) [13] method for comparison. We used the implementation in MINC toolkit (https://bic-mni.github.io/) with the default parameters, including a patch radius of two voxels and a search radius of five voxels. All images were preprocessed in the same way as described in section II-F, but with the addition of a linear intensity normalization step mapping all voxel intensities into the range [0, 100]. Mean Dice coefficients for each source → target task are reported in Table III. Compared to the CNN-based methods, PBS was found to be more sensitive to disparities between training and testing domains. Over all structures and inter-domain tasks, PBS produced a very low mean Dice coefficient of 64.4% across all structures and all source → target tasks. On the other hand, when training on the target domains using a 5-fold cross-validation, PBS markedly improved (mean Dice coefficient of 81.7%), though its performance was still worse compared to the CNN-based method both with (mean Dice coefficient of 83.9%) and without (mean Dice coefficient of 82.7%) data augmentation. The observed disparity in the performance of PBS between inter- and intra-domain applications can be understood in light of the following considerations. First, since PBS relies on the *L*_2_ distance to estimate patch similarities, accurate label propagation requires that tissues have similar intensity values across training and testing images. Second, since PBS only extracts candidate patches for label fusion within local search windows, it requires excellent spatial alignment between training and target images. In general, achieving sufficiently consistent intensity normalization and spatial alignment across images from different domains is a highly challenging problem; while using labels (e.g. tissue maps) can help intensity normalization and registration achieve more consistent results across domains, this requires segmentation, which, as highlighted in the previous results, is particularly burdened by performance degradation when applied across domains.

**TABLE 3.** Patch-based segmentation applied to all source → target tasks. Mean Dice coefficients (standard deviation in parentheses) are reported for each source → target task over all structures. The bottom row shows the performance obtained by a 5-fold cross validation on the target domain. A: ADNI, C: CANDI, O: OASIS.

| | A → C | A → O | C → A | C → O | O → A | O → C | All |
| --- | --- | --- | --- | --- | --- | --- | --- |
| Source | 63.6 (1.5) | 74.7 (1.0) | 53.4 (1.7) | 68.4 (1.4) | 65.1 (1.7) | 66.8 (1.4) | 64.4 (1.7) |
| Target | 82.6 (0.8) | 83.1 (0.6) | 80.5 (0.8) | 83.1 (0.6) | 80.5 (0.8) | 82.6 (0.8) | 81.7 (0.7) |

### 3.4 Adaptation visualization

To visualize the effect of data augmentation and explicit domain adaptation on the trained networks, we used the t-SNE algorithm [33] to reduce the dimensionality of sample sets of features produced by the various networks. In Fig. 8, we consider the CANDI → ADNI adaptation, and plot sample feature sets (here, feature maps extracted immediately prior to the final softmax layer) produced by 1000 uniformly sampled input patches from each domain. The unadapted network without data augmentation produced highly disparate, with samples from the target domain tending to be highly concentrated in the center of the source domain feature distribution. The addition of data augmentation tended to slightly disperse the target domain samples away from the center, while the combination of data augmentation with both self-ensembling and the domain-adverserial method produced more consistent feature distributions across domains.

**Fig. 8.**
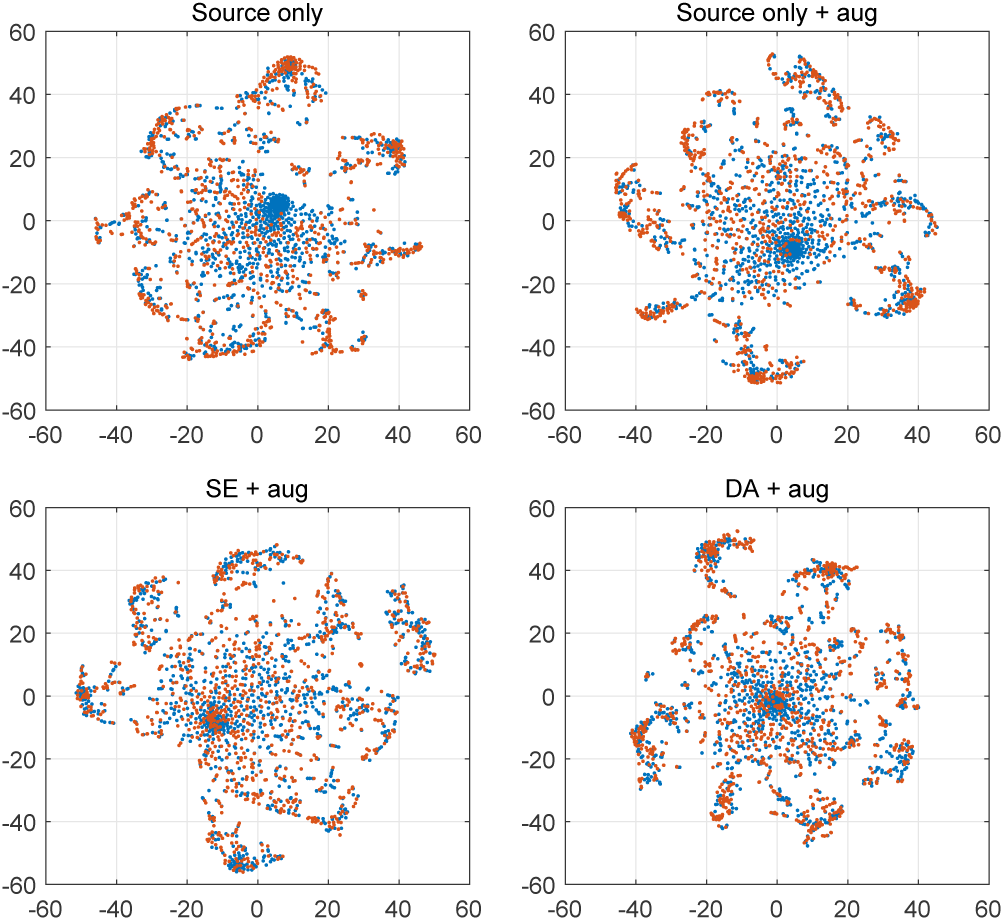
Low-dimensional embedding of network-extracted features from unadapted and adapted networks in the CANDI → ADNI task. Uniformly sampled patches from the ADNI (blue) and CANDI (orange) datasets were passed through the unadapted networks with no data augmentation (top left), with data augmentation (top right), and data augmentation combined with self-ensembling (bottom left) and the domain-adverarial method (bottom right).

## 4 Discussion

Despite the superficial similarity between T1w datasets, domain adaptation is a highly challenging task due to an abundance of low-level (e.g. brightness, contrast, noise and resolution) differences caused by varying pulse sequence parameters and scanning hardware, in addition to high-level anatomical differences due to the age and health/disease of the imaged demographic. In this work we have developed and validated a novel CNN-based method for fully unsupervised domain adaptation in automated neuroanatomical segmentation of T1w MRI. The approach is fully unsupervised on the target domain and does not require additional domain-specific training data, allowing users to easily and more effectively process their data of interest using prelabelled images from an arbitrary domain. This is particularly important for processing modern large-scale conglomerate datasets consisting of images from various centres, as well a necessary quality before such automated methods can be confidently applied in clinical environments. Combining an extensive data augmentation scheme (designed to mimic inter-domain variability in T1w MRI) with a novel self-ensembling approach, our proposed method demonstrated increased performance compared to the baseline network trained on the source domain only, a previously published domain-adversarial method for domain adaptation [7], and a classical patch-based segmentation method [13]. Considering the performance of the baseline network (mean Dice coefficient of 73.2% across all structures and all source → target adaptations), our fully unsupervised method for domain adaptation improved performance by 5.2%, closing the gap by 49% relative to fully supervised training on the target domain with data augmentation (mean Dice coefficient of 83.9%), and generally avoided the more serious segmentation errors produced by the baseline network.

Rather than using hand-crafted data augmentation transformations, it is also possible to learn transformations which map the style (low-level appearance) of images across domains [34, 35, 36]. One difficulty with this approach is to ensure that the learned transformations sufficiently translate style while (1) remaining realistic, and (2) preserving content (anatomy) such that they are label-preserving [37]. As noted by Cohen et al. [38], this is particularly problematic when using approaches based on adversarial losses (e.g. CycleGAN [34]) which aim to match the translation output with the distribution of the target domain, commonly introducing anatomical artifacts into the translated output, and introducing inconsistencies between the input labels and the transformed outputs. This could possibly be remedied by attaching further constraints to the learned mapping, e.g. by requiring that the correlation between input and style-transferred outputs be maximized. A second difficulty is that style-transfer mappings are generally not stochastic [39] as required for compatibility with self-ensembling. Thus, the combination of learned style-transfer mappings with the random data augmentations used in this work may result in further performance gains.

Semi-supervised approaches for domain adaptation can also be considered. This class of methods requires additional but limited quantities of labelled data from the target domain, which can be used, for example, for fine-tuning the network parameters (also called ‘transfer learning’) (see [6, 40] for an example of transfer learning applied to segmentation in MRI). While less practical, semi-supervised approaches work orthogonally to unsupervised approaches. Therefore, the combination of unsupervised (using all unlabelled target domain data) and semi-supervised (using a small quantity of labelled target domain data) approaches may provide further performance gains. Active learning approaches [41] could also be used to reduce the amount of required manual effort, e.g. by requiring labels on a smaller subset of the most informative samples on the target domain.

Finally, this work concerns pairwise adaptation from a single source domain to a single target domain. In practice, users may have access to pre-labelled training images from multiple source domains. In this case, applying pairwise adaptation approaches may be suboptimal, as they fail to leverage the shared information across domains. A natural solution would be to pool all available training data together and then proceed using a pairwise adaptation approach. However, some source domains may not be useful for adaptation to particular domains, and this approach may in certain cases actually hurt performance [42]. The question of how to optimally leverage multiple source domains for adaptation is indeed an active field of research in the wider computer vision literature. For example, Duan et al. [43] propose a general method where networks trained from each source can be weighted and then combined to make a final decision on the target domain. Alternatively, explicit multi-source domain adaptation models can be constructed, such as in the work of Zhao et al. [44], which extends the domain-adversarial adaptation method to multiple source domains by back-propagating gradients from multiple source domains in proportion to their similarity to the target domain. Developing and extending these and similar methods specifically for the problem of domain-adaptation in MR segmentation is a promising direction for future work.

